# Revisiting the Origin of the Octoploid Strawberry

**DOI:** 10.1101/665216

**Authors:** Aaron Liston, Na Wei, Jacob Tennessen, Junmin Li, Ming Dong, Tia-Lynn Ashman

## Abstract

The cultivated strawberry, *Fragaria ×ananassa*, originated in France approximately 270 years ago via hybridization between two wild species introduced from North and South America. Both the cultivated strawberry and its parental species are octoploids with 2*n*=*8x*=56 chromosomes. In the recent publication of the genome of the cultivated strawberry, the authors present a novel phylogenetic hypothesis, proposing that each of the four subgenomes originated from a different 2*n*=2*x*=14 diploid progenitor. They further suggest that the hexaploid species *Fragaria moschata* was a direct ancestor of the strawberries. We reanalyzed the four octoploid subgenomes in a phylogenomic context, and found that only two extant diploids were progenitors, a result that is consistent with several previous studies. We also conducted a phylogenetic analysis of genetic linkage-mapped loci in the hexaploid *F. moschata*, and resolved its origin as independent of the octoploids. We identified assumptions in their tree-searching algorithm that prevented it from accepting extinct or unsampled progenitors, and we argue that this is a critical weakness of their approach. Correctly identifying their diploid progenitors is important for understanding and predicting the responses of polyploid plants to climate change and associated environmental stress.

## Introduction

The publication by Edger et al.^1^ of the first chromosome-scale genome assembly of the octoploid strawberry *Fragaria ×ananassa* cultivar ‘Camarosa’ represents a significant scientific advance and provides a foundational resource for this important cultivated plant. The authors also identified four diploid species of *Fragaria* as the progenitors of the wild octoploid species of strawberry (Table 1, Fig. 1a). Three of these species are distributed in Asia, the fourth in western North America. Edger et al.^1^ further hypothesized that the hexaploid species *F. moschata* “may be evolutionary intermediate between the diploids and wild octoploid species”. This predicts that the hexaploid’s three subgenomes should correspond to these three Asian diploid ancestors (Fig. 1a).

**Table 1:** Summary of recent hypotheses for the diploid ancestors of the wild octoploid strawberries, *Fragaria virginiana* and *F. chiloensis*. These two species were introduced to Europe and hybridized approximately 270 years ago, creating *F. ×ananassa*, the cultivated strawberry (reviewed in ref. ^5^). Phylogenetic analyses support a single origin between 0.43-1.62 million years ago^9^. Kamneva et al.^3^ found no evidence for additional progenitors beyond *F. vesca* and *F. iinumae*, and while their results were consistent with 3 *F. iinumae*-like subgenomes, this could not be directly evaluated. Yang and Davis^4^ proposed an additional unknown progenitor, and questioned whether each subgenome had a single diploid ancestry. The A,B,C,D genome designations are to provide a common reference, and were not used in these studies.

| genome A | genome B | genome C | genome D | authors |
| --- | --- | --- | --- | --- |
| *vesca*<br>*bracteata* | *iinumae* | *iinumae*-<br>related | *iinumae*-<br>related | This study |
| *vesca*<br>*bracteata* | *iinumae* | *nipponica* | *viridis* | Edger et al. 2019 |
| *vesca* | *iinumae* | - | - | Kamneva et al. 2017 |
| *vesca* | *iinumae* | *bucharica* | *viridis* | Yang & Davis 2017 |
| *vesca*<br>*bracteata* | *iinumae* | *iinumae*-<br>related | *iinumae*-<br>related | Tennessen et al. 2014 |

**Figure 1.**
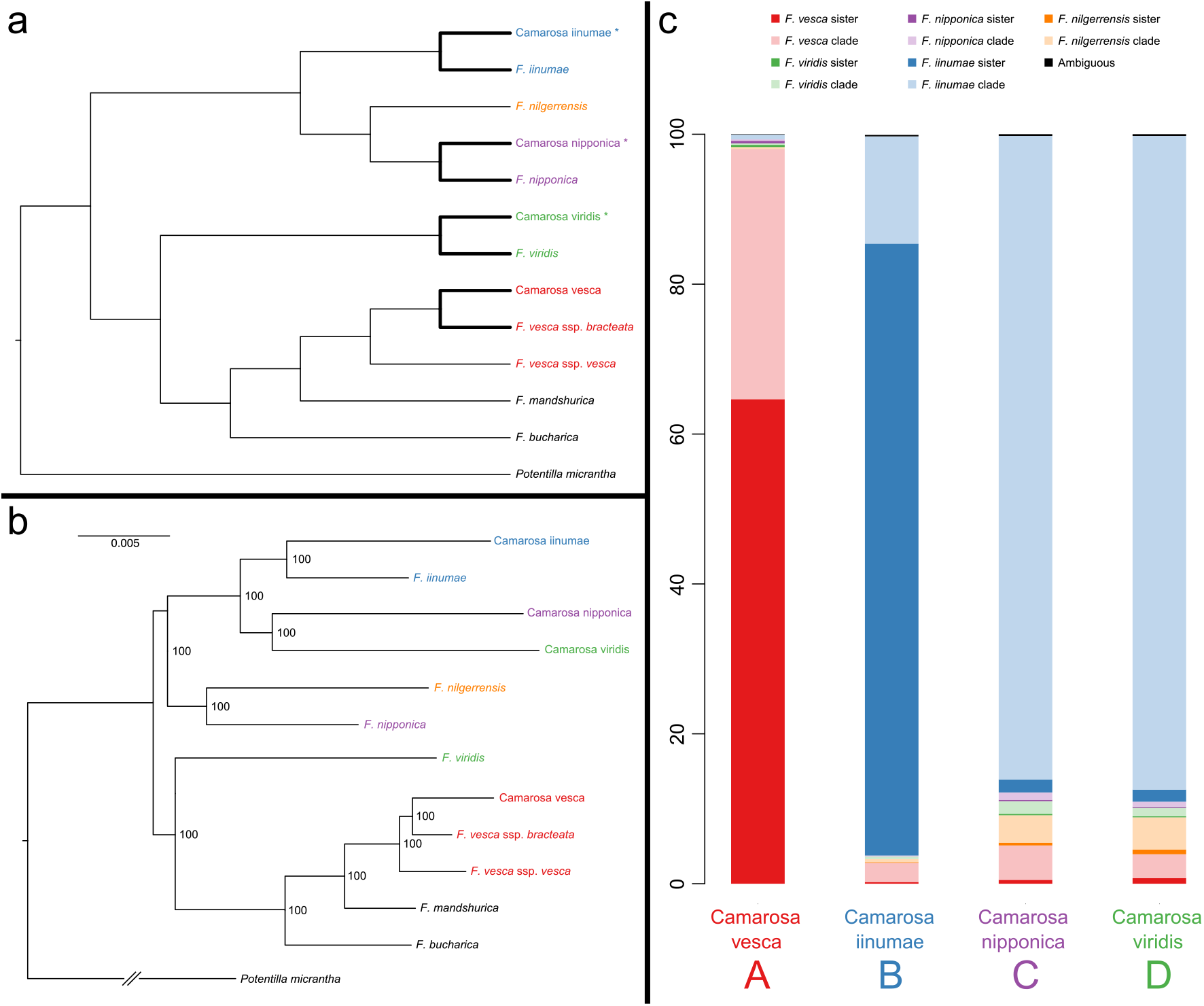
Phylogenomics of octoploid subgenomes. **a**, The phylogenetic hypothesis presented by Edger et al.^1^ suggesting that each octoploid subgenome has a different extant diploid progenitor, and that three of these subgenomes (denoted by asterisks) comprise the hexaploid species *F. moschata*. The bold lines represent the topology constraint we applied in the Shimodaira-Hasegawa test of the Edger et al.^1^ hypothesis vs. our phylogenomic results (Table 3). **b**, Maximum likelihood estimate of phylogeny for base chromosome 1, all nodes have 100% bootstrap support. This topology is shared by five of the seven chromosomes (Fig 1). **c**, Summary of the phylogenetic positions of the four octoploid subgenomes in maximum likelihood estimates of phylogeny in 2191 non-overlapping windows of 100 kb across the 7 base chromosomes. The octoploid subgenomes A ‘Camarosa vesca’ and B ‘Camarosa iinumae’ generally are sister to their eponymous diploids *F. vesca* (red) and *F. iinumae* (blue). In contrast, subgenome C ‘Camarosa nipponica’ and subgenome D ‘Camarosa viridis’, predominantly share a most recent common ancestor (clade) with *F. iinumae* (light blue). See Fig. 3 for an explanation of the sister vs. clade categorization and the results for all 2191 windows. For data sources and phylogenetic methods, see Materials and Methods.

## Results and Discussion

Since these phylogenetic conclusions conflict with several recent studies^2–4^ (Table 1), we conducted two new analyses: a chromosome-scale phylogenomic analysis of the four *F. ×ananassa* subgenomes (Fig. 1b,c; Table 2, Figs. 2–3) and a phylogenetic analysis of genetic linkage-mapped loci in *F. moschata* (Figs. 4–5). We evaluated the phylogenetic hypotheses of Edger et al.^1^ using topology tests and found no evidence of *F. nipponica* and *F. viridis* ancestry in the octoploid (Table 3) and supported independent origins of the hexaploid and octoploid strawberries (Table 4).

**Table 2:**
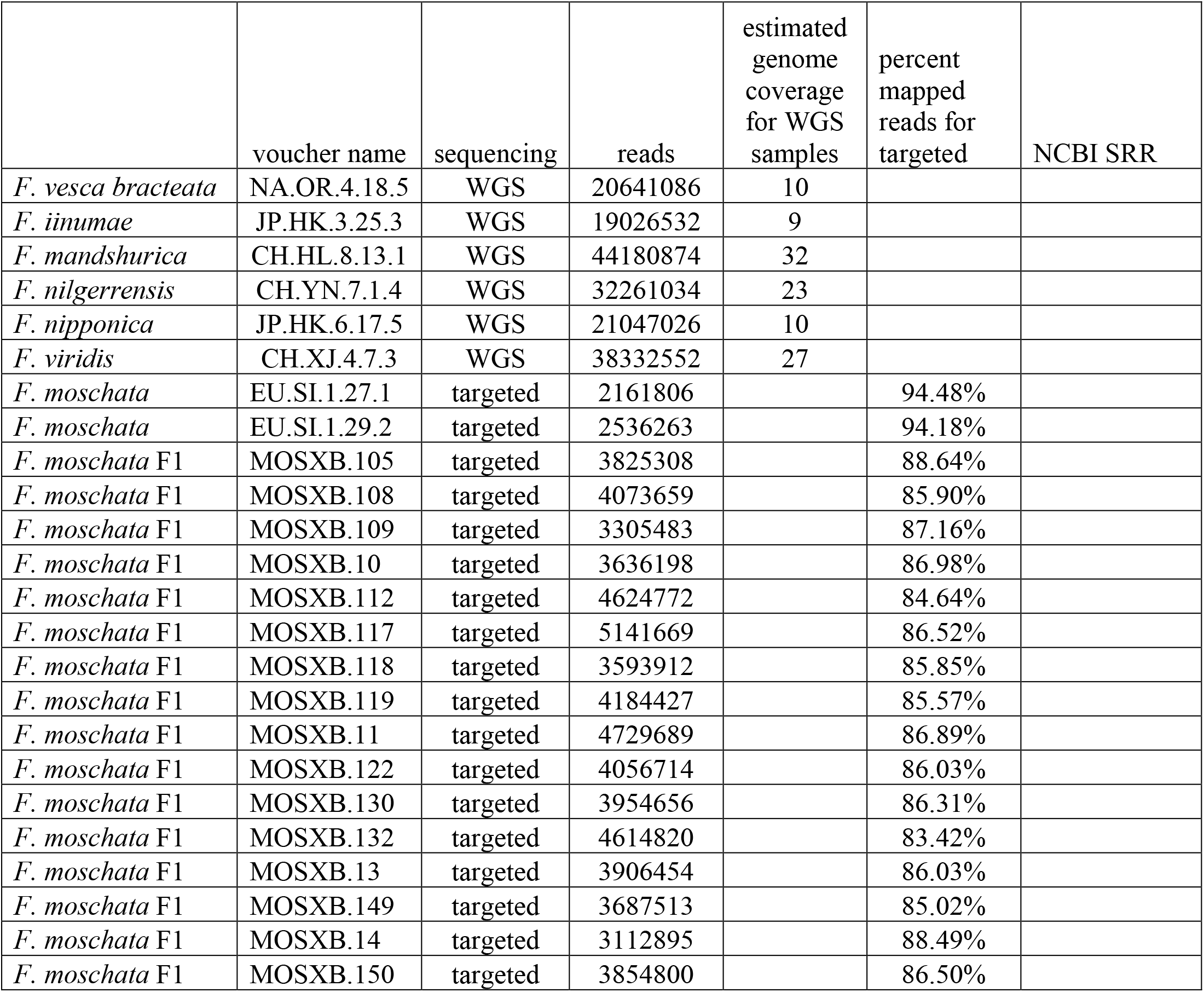
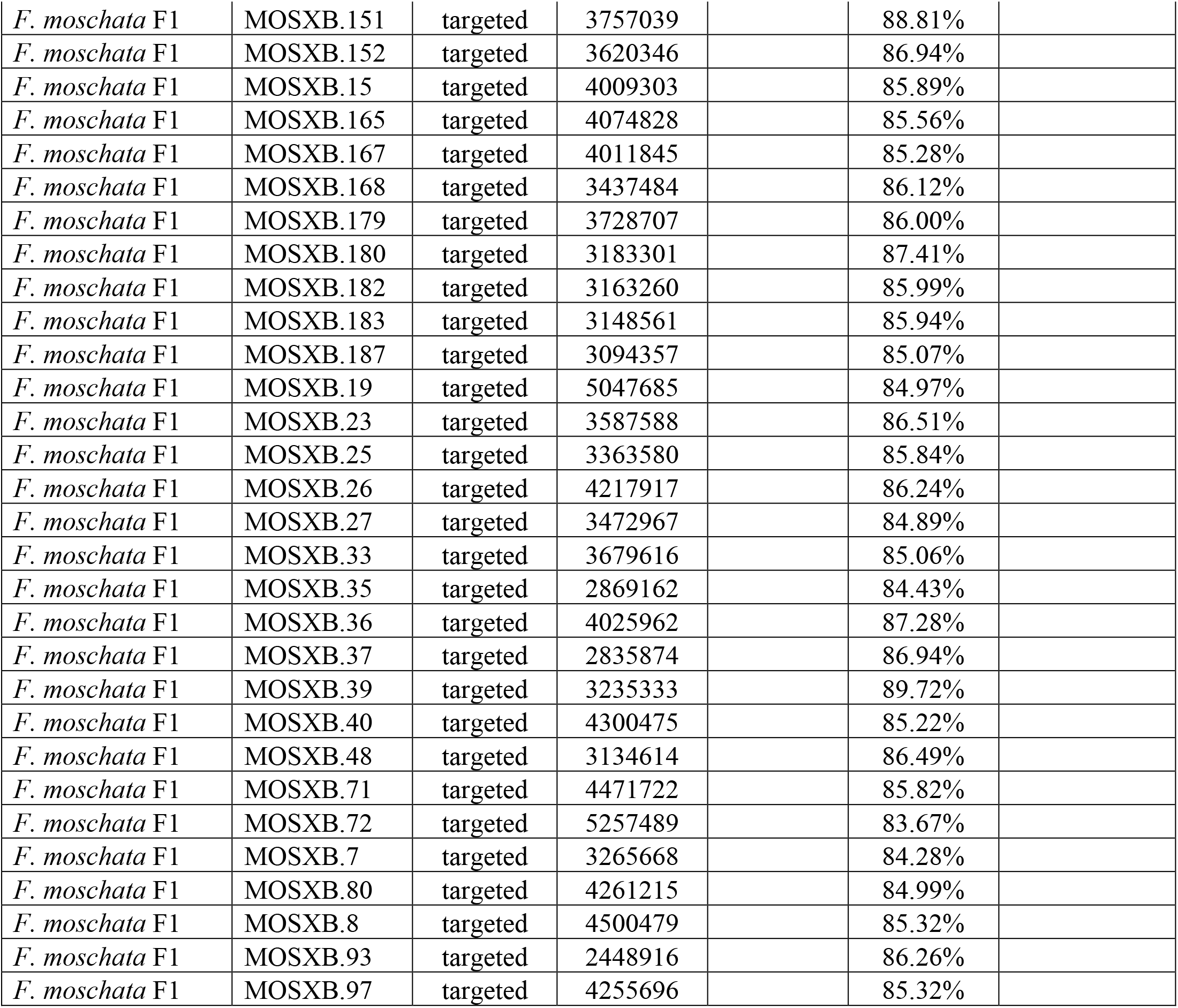
New sequences reported here. Detailed locality data is available at http://wildstrawberry.org

|  | voucher name | sequencing | reads | estimated<br>genome<br>coverage<br>for WGS<br>samples | percent<br>mapped<br>reads for<br>targeted | NCBI SRR |
| --- | --- | --- | --- | --- | --- | --- |
| *F. vesca bracteata* | NA.OR.4.18.5 | WGS | 20641086 | 10 |  |  |
| *F. iinumae* | JP.HK.3.25.3 | WGS | 19026532 | 9 |  |  |
| *F. mandshurica* | CH.HL.8.13.1 | WGS | 44180874 | 32 |  |  |
| *F. nilgerrensis* | CH.YN.7.1.4 | WGS | 32261034 | 23 |  |  |
| *F. nipponica* | JP.HK.6.17.5 | WGS | 21047026 | 10 |  |  |
| *F. viridis* | CH.XJ.4.7.3 | WGS | 38332552 | 27 |  |  |
| *F. moschata* | EU.SI.1.27.1 | targeted | 2161806 |  | 94.48% |  |
| *F. moschata* | EU.SI.1.29.2 | targeted | 2536263 |  | 94.18% |  |
| *F. moschata* F1 | MOSXB.105 | targeted | 3825308 |  | 88.64% |  |
| *F. moschata* F1 | MOSXB.108 | targeted | 4073659 |  | 85.90% |  |
| *F. moschata* F1 | MOSXB.109 | targeted | 3305483 |  | 87.16% |  |
| *F. moschata* F1 | MOSXB.10 | targeted | 3636198 |  | 86.98% |  |
| *F. moschata* F1 | MOSXB.112 | targeted | 4624772 |  | 84.64% |  |
| *F. moschata* F1 | MOSXB.117 | targeted | 5141669 |  | 86.52% |  |
| *F. moschata* F1 | MOSXB.118 | targeted | 3593912 |  | 85.85% |  |
| *F. moschata* F1 | MOSXB.119 | targeted | 4184427 |  | 85.57% |  |
| *F. moschata* F1 | MOSXB.11 | targeted | 4729689 |  | 86.89% |  |
| *F. moschata* F1 | MOSXB.122 | targeted | 4056714 |  | 86.03% |  |
| *F. moschata* F1 | MOSXB.130 | targeted | 3954656 |  | 86.31% |  |
| *F. moschata* F1 | MOSXB.132 | targeted | 4614820 |  | 83.42% |  |
| *F. moschata* F1 | MOSXB.13 | targeted | 3906454 |  | 86.03% |  |
| *F. moschata* F1 | MOSXB.149 | targeted | 3687513 |  | 85.02% |  |
| *F. moschata* F1 | MOSXB.14 | targeted | 3112895 |  | 88.49% |  |
| *F. moschata* F1 | MOSXB.150 | targeted | 3854800 |  | 86.50% |  |
| *F. moschata* F1 | MOSXB.151 | targeted | 3757039 |  | 88.81% |  |
| *F. moschata* F1 | MOSXB.152 | targeted | 3620346 |  | 86.94% |  |
| *F. moschata* F1 | MOSXB.15 | targeted | 4009303 |  | 85.89% |  |
| *F. moschata* F1 | MOSXB.165 | targeted | 4074828 |  | 85.56% |  |
| *F. moschata* F1 | MOSXB.167 | targeted | 4011845 |  | 85.28% |  |
| *F. moschata* F1 | MOSXB.168 | targeted | 3437484 |  | 86.12% |  |
| *F. moschata* F1 | MOSXB.179 | targeted | 3728707 |  | 86.00% |  |
| *F. moschata* F1 | MOSXB.180 | targeted | 3183301 |  | 87.41% |  |
| *F. moschata* F1 | MOSXB.182 | targeted | 3163260 |  | 85.99% |  |
| *F. moschata* F1 | MOSXB.183 | targeted | 3148561 |  | 85.94% |  |
| *F. moschata* F1 | MOSXB.187 | targeted | 3094357 |  | 85.07% |  |
| *F. moschata* F1 | MOSXB.19 | targeted | 5047685 |  | 84.97% |  |
| *F. moschata* F1 | MOSXB.23 | targeted | 3587588 |  | 86.51% |  |
| *F. moschata* F1 | MOSXB.25 | targeted | 3363580 |  | 85.84% |  |
| *F. moschata* F1 | MOSXB.26 | targeted | 4217917 |  | 86.24% |  |
| *F. moschata* F1 | MOSXB.27 | targeted | 3472967 |  | 84.89% |  |
| *F. moschata* F1 | MOSXB.33 | targeted | 3679616 |  | 85.06% |  |
| *F. moschata* F1 | MOSXB.35 | targeted | 2869162 |  | 84.43% |  |
| *F. moschata* F1 | MOSXB.36 | targeted | 4025962 |  | 87.28% |  |
| *F. moschata* F1 | MOSXB.37 | targeted | 2835874 |  | 86.94% |  |
| *F. moschata* F1 | MOSXB.39 | targeted | 3235333 |  | 89.72% |  |
| *F. moschata* F1 | MOSXB.40 | targeted | 4300475 |  | 85.22% |  |
| *F. moschata* F1 | MOSXB.48 | targeted | 3134614 |  | 86.49% |  |
| *F. moschata* F1 | MOSXB.71 | targeted | 4471722 |  | 85.82% |  |
| *F. moschata* F1 | MOSXB.72 | targeted | 5257489 |  | 83.67% |  |
| *F. moschata* F1 | MOSXB.7 | targeted | 3265668 |  | 84.28% |  |
| *F. moschata* F1 | MOSXB.80 | targeted | 4261215 |  | 84.99% |  |
| *F. moschata* F1 | MOSXB.8 | targeted | 4500479 |  | 85.32% |  |
| *F. moschata* F1 | MOSXB.93 | targeted | 2448916 |  | 86.26% |  |
| *F. moschata* F1 | MOSXB.97 | targeted | 4255696 |  | 85.32% |  |

**Figure 2:**
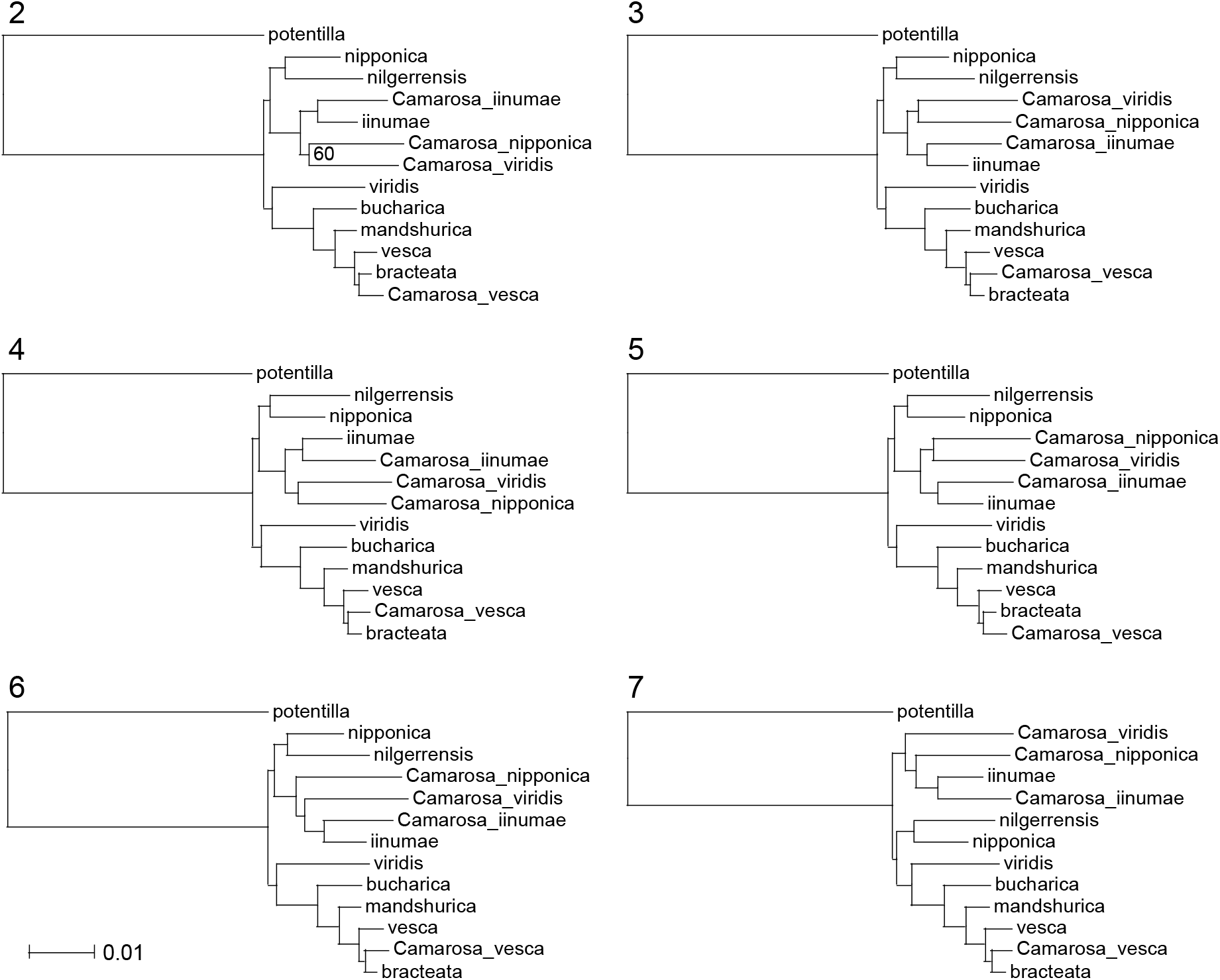
Maximum likelihood estimate of phylogeny for base chromosomes 2-7. Bootstrap values for all nodes are 100%, unless indicated.

**Figure 3:**
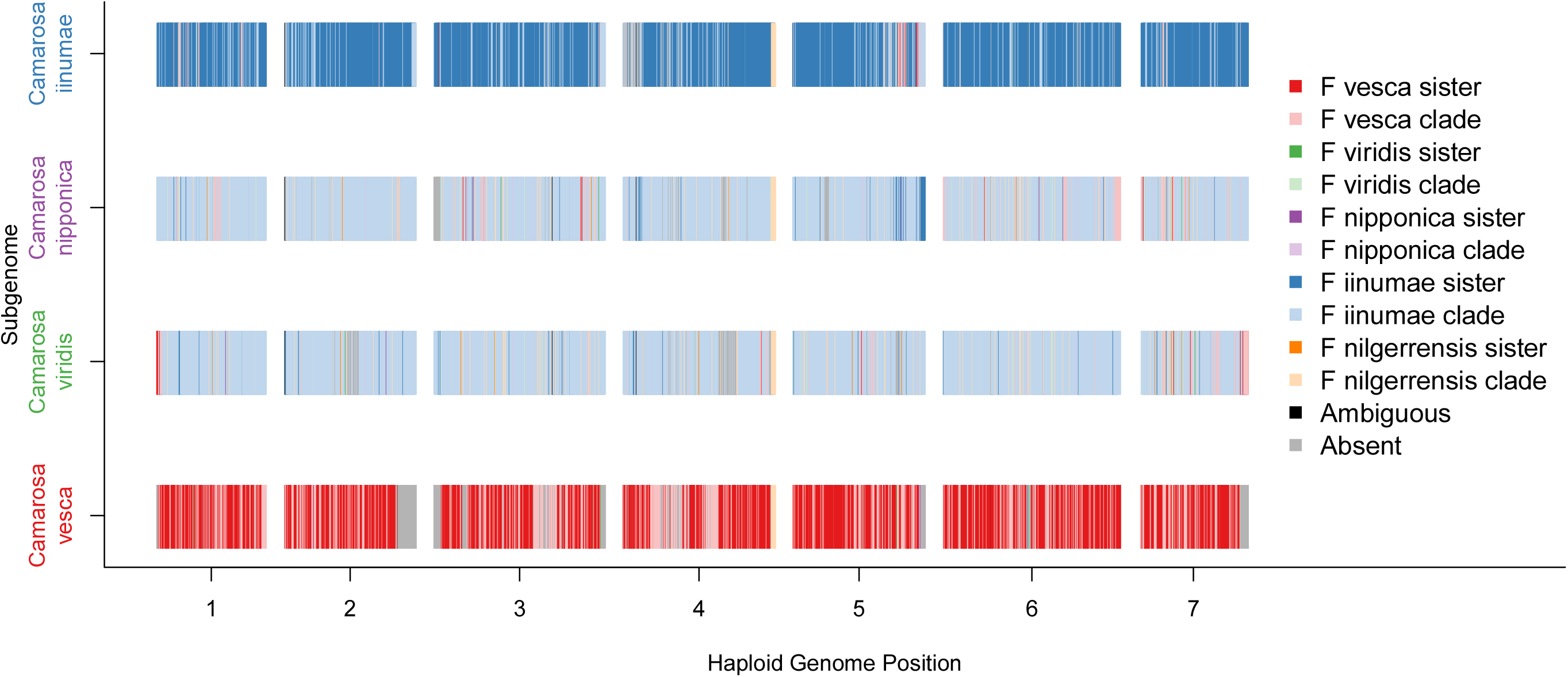
Phylogenetic positions of the four octoploid subgenomes in maximum likelihood estimates of phylogeny in 2191 non-overlapping windows of 100 kb across the 7 base chromosomes. Phylogenetic matrices averaged 89077 nucleotides per 100 kb window (Supplementary Table 1). For each window, we recorded the diploid species sharing the most recent common ancestor (MRCA) with each octoploid subgenome, and we further noted when this diploid was “sister” to that subgenome. If the MRCA diploid was not sister to that subgenome, we labelled these as “clade”. This generally occurred when subgenomes were sister to each other, e.g. ‘Camarosa viridis’ and ‘Camarosa nipponica’ in Fig. 1b. The two subspecies of *F. vesca* and the closely related *F. mandshurica* and *F. bucharica* were grouped together for MRCA scoring, and combined into a single color category. We restricted the category “vesca sister” to windows where the octoploid subgenome is sister to one subspecies of *F. vesca*; all other positions were scored as “vesca clade”. More than one diploid species shared a MRCA with multiple subgenomes in 0.2% of the 2191 trees, and these were labelled “ambiguous”. When an octoploid subgenome was excluded from a 100 kbp window (<10% of aligned sites) it was scored “absent”. These regions primarily correspond to large duplications, e.g. at the 3’ end of chromosome 2 and 7 in the *F. vesca* subgenome.

**Figure 4:**
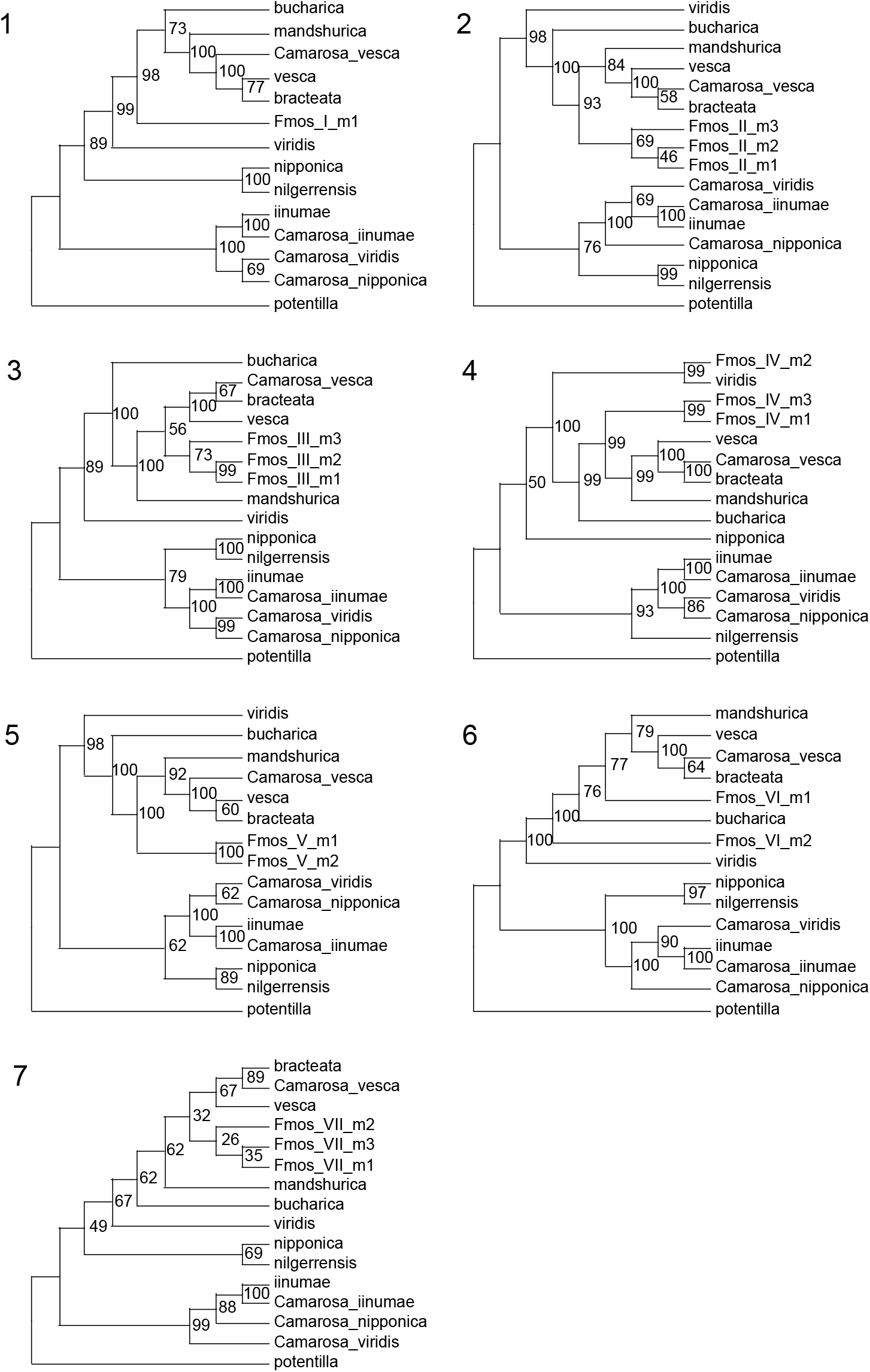
Maximum-likelihood phylogenies of base chromosomes 1–7 involving the maternal linkage groups (LGs) of the hexaploid *Fragaria moschata*. These LGs are named after species (‘Fmos’), chromosome (I–VII), maternal map (m) and LG number (1–3). Some chromosomes (e.g. 1, 5 and 6) have incomplete *F. moschata* LG numbers. Numbers associated with branches are ML bootstrap support values from 100 replicates.

**Figure 5:**
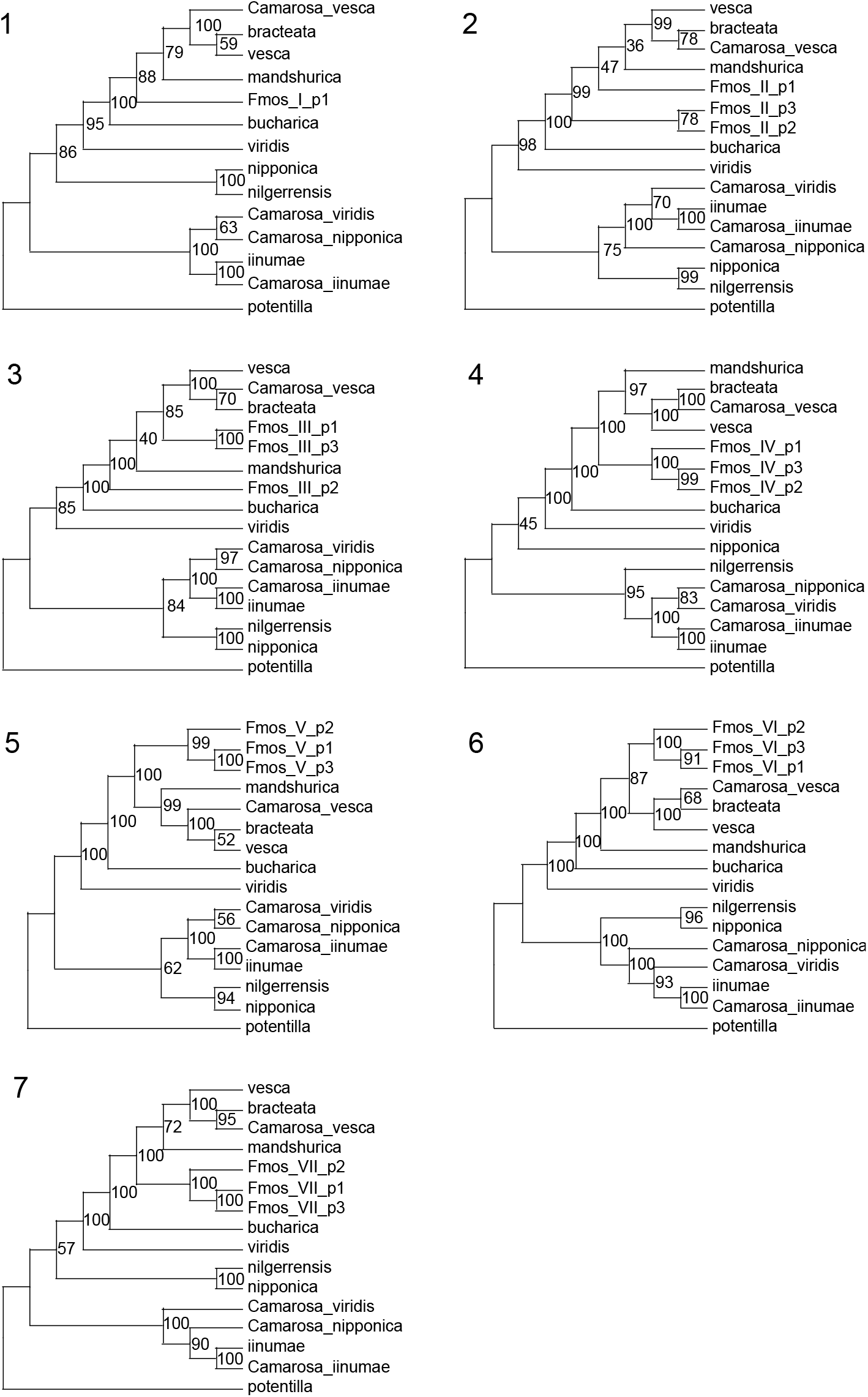
Maximum-likelihood phylogenies of base chromosomes 1–7 involving the paternal linkage groups (LGs) of the hexaploid *Fragaria moschata*. These LGs are named after species (‘Fmos’), chromosome (I–VII), paternal map (p) and LG number (1–3). Chromosome 1 has incomplete *F. moschata* LG numbers. Numbers associated with branches are ML bootstrap support values from 100 replicates.

**Table 3:** Shimodaira–Hasegawa (SH) tests support an alternative scenario of the evolutionary origins of the octoploid strawberry, relative to the hypothesis of Edger *et al*. (2019).

|  | Chromosome | 1 | 2 | 3 | 4 | 5 | 6 | 7 |
| --- | --- | --- | --- | --- | --- | --- | --- | --- |
| ML unconstrained tree | -Log<sub>e</sub>L | 43776167 | 53635963 | 67303204 | 58266330 | 53739632 | 72891475 | 43776415 |
| Edger *et al.* (2019) hypothesis |  |  |  |  |  |  |  |  |
| (*Camarosa_vesca*,*bracteata*),<br>(*Camarosa_iinumae*,*iinumae*),<br>(*Camarosa_viridis*,*viridis*),<br>(*Camarosa_nipponica*,*nipponica*) | Δ -Log<sub>e</sub>L | 158226** | 219842** | 198938** | 190332** | 192305** | 245147** | 114341** |

-Log<sub>e</sub>L: negative log likelihood; Δ -Log<sub>e</sub>L: the difference in -Log<sub>e</sub>L between the best ML constraint tree and the ML unconstrained tree for each of chromosomes 1–7.
The *P* values of the SH tests are denoted as asterisks: \*\*, *P* < 0.01.

**Table 4:** Shimodaira–Hasegawa (SH) tests do not support the hypothesis that the hexaploid *Fragaria moschata* was the evolutionary intermediate of the octoploid strawberry, as evidenced by moschata maternal LGs (m1-m3) and paternal LGs (p1-p3).

|  | Chromo<br>some | 1 | 2 | 3 | 4 | 5 | 6 | 7 |
| --- | --- | --- | --- | --- | --- | --- | --- | --- |
| **Maternal** |  |  |  |  |  |  |  |  |
| ML unconstrained tree | -Log<sub>e</sub>L | 89129 | 278511 | 208456 | 213813 | 206867 | 392846 | 215159 |
| Constrained trees |  |  |  |  |  |  |  |  |
| ((Camarosa_viridis,m1),(Camarosa_iinumae,m2),(Camarosa_nipponica,m3)); | Δ -Log<sub>e</sub>L | - | 831** | 685** | 558** | 341** | 761** | 104** |
| ((Camarosa_viridis,m1),(Camarosa_iinumae,m3),(Camarosa_nipponica,m2)); | Δ -Log<sub>e</sub>L | - | 846** | 381** | 559** | 341** | 751** | 100** |
| ((Camarosa_viridis,m2),(Camarosa_iinumae,m1),(Camarosa_nipponica,m3)); | Δ -Log<sub>e</sub>L | - | 831** | 715** | 543** | 337** | 809** | 134** |
| ((Camarosa_viridis,m2),(Camarosa_iinumae,m3),(Camarosa_nipponica,m1)); | Δ -Log<sub>e</sub>L | - | 846** | 715** | 567** | 382** | 809** | 425** |
| ((Camarosa_viridis,m3),(Camarosa_iinumae,m1),(Camarosa_nipponica,m2)); | Δ -Log<sub>e</sub>L | - | 928** | 643** | 544** | 337** | 750** | 131** |
| ((Camarosa_viridis,m3),(Camarosa_iinumae,m2),(Camarosa_nipponica,m1)); | Δ -Log<sub>e</sub>L | - | 922** | 684** | 568** | 382** | 760** | 153** |

| **Paternal** |  |  |  |  |  |  |  |  |
| --- | --- | --- | --- | --- | --- | --- | --- | --- |
| ML unconstrained tree | -Log<sub>e</sub>*L* | 87227 | 275286 | 211795 | 215665 | 208624 | 382540 | 221318 |
| Constrained trees |  |  |  |  |  |  |  |  |
| ((Camarosa_viridis,p1),(Camarosa_iinumae,p2),(Camarosa_nipponica,p3)); | Δ -Log<sub>e</sub>*L* | - | 1033** | 1024** | 1461** | 1112** | 1542** | 1113** |
| ((Camarosa_viridis,p1),(Camarosa_iinumae,p3),(Camarosa_nipponica,p2)); | Δ -Log<sub>e</sub>*L* | - | 1046** | 1028** | 1290** | 1112** | 1522** | 882** |
| ((Camarosa_viridis,p2),(Camarosa_iinumae,p1),(Camarosa_nipponica,p3)); | Δ -Log<sub>e</sub>*L* | - | 1072** | 1172** | 1478** | 1114** | 1562** | 1136** |
| ((Camarosa_viridis,p2),(Camarosa_iinumae,p3),(Camarosa_nipponica,p1)); | Δ -Log<sub>e</sub>*L* | - | 1068** | 1176** | 1290** | 1141** | 1547** | 884** |
| ((Camarosa_viridis,p3),(Camarosa_iinumae,p1),(Camarosa_nipponica,p2)); | Δ -Log<sub>e</sub>*L* | - | 1046** | 1013** | 1478** | 755** | 1504** | 860** |
| ((Camarosa_viridis,p3),(Camarosa_iinumae,p2),(Camarosa_nipponica,p1)); | Δ -Log<sub>e</sub>*L* | - | 1030** | 1041** | 1461** | 755** | 1528** | 862** |

We believe that the “phylogenetic analysis of subgenomes” tree-searching algorithm (PhyDS) developed by Edger et al.^1^ was responsible for the unsupported identification of *F. nipponica* and *F. viridis* ancestry for the octoploids. Their analysis was based on 8,405 individual gene trees compiled from the annotation of the newly assembled octoploid genome and 31 transcriptomes from 12 diploid *Fragaria* species. In each gene tree, when multiple octoploid genes resolved as sister to the same diploid, these were treated as in-paralogs, and ignored. This forces each of the four octoploid subgenomes to have a different diploid ancestor, an assumption that is at odds with most classical genetic hypotheses for the octoploid ancestry (reviewed in ref.^5^) as well as the results of molecular phylogenetic analyses of *Fragaria*^2–4^.

In effect, Edger et al.^1^ only considered gene trees as informative when genes from a single octoploid and diploid comprise an exclusive clade. This is an effective strategy for subgenomes A and B (Table 1), where a high proportion of PhyDS resolved gene trees (their Supplementary Table 8) and 100-kb windows (labelled “sister” in Fig. 1c) resolve these with *F. vesca* and *F. iinumae*, respectively. However, the approach fails for subgenomes C and D (Table 1) where a smaller percentage of PhyDS accepted gene trees (their Supplementary Table 8), and very few 100-kb windows (Fig. 1c), resolve an exclusive clade relationship between these subgenomes and *F. nipponica* and *F. viridis*, respectively.

The rationale of Edger et al.^1^ for treating in-paralogs in this way was to avoid errors when homeologous exchange has “replaced” the syntelog of one subgenome with another, as illustrated in their Supplementary Figure 8c. According to their Supplementary Table 10, 11.4% of the genome has experienced homeologous exchange, suggesting that 90% of the syntelog gene trees should correctly resolve subgenome ancestry. In contrast, only ca. 3% of syntelogs across the genome resulted in a subgenome assignment by PhyDS. This indicates that their in-paralog exclusion criterion was too strict. Our results suggest that the great majority of trees rejected by PhyDS resolved *F. iinumae* as the diploid progenitor of three subgenomes (Fig. 1b, Fig. 2, Supplementary Table 2). This topology matches the examples of “incorrectly identified as progenitor” in their Supplementary Figure 8c, but in our view reflects the most likely evolutionary scenario for the origin of the octoploid strawberries.

Our phylogenomic approach found that 12.5% of the genome has experience homeologous exchange (Fig. 1c, Table 2), similar to the Edger et al. estimate of 11.4%. Although none of the 2191 trees from 100-kb windows across the genome (Table 4) match the Edger et al. hypothesis (Fig. 1a), a small number do resolve subgenome C or subgenome D with the diploid species *F. nipponica* and *F. viridis*, respectively (Fig. 1c, Supplementary Table 2). However, these are similar to the number of trees that resolve the opposite species, or *F. nilgerrensis*, as the most closely related diploid. We suspect that these results can be attributed to incomplete lineage sorting.

In 81.4% of the 2011 trees where subgenome A (Camarosa vesca) was resolved with the *F. vesca* clade, its most recent common ancestor or sister taxon was *F. vesca* subsp. *bracteata* (Supplementary Table 2), native to northwest North America. This is consistent with the PhyDS results of Edger et al.^1^, and most previous studies^2–4,6,7^. Likewise, our results and previous studies resolve subgenome B (Camarosa iinumae) with *F. iinumae* from Japan. Subgenomes C and D are most commonly resolved as sister to each other (Fig. 1b, Fig. 2), and may represent an autotetraploid ancestor^2^, although this requires cytogenetic confirmation. *Fragaria iinumae* is the closest diploid progenitor of these two subgenomes (Fig. 1c, Fig. 3). Whether they originated with an extinct species^8^ or an unsampled population of *F. iinumae* (e.g. from Sakhalin Island) remains to be determined. The origin of the wild octoploid strawberry species likely occurred in Pleistocene Beringia, which is temporally consistent with the estimated age of the octoploids^6,9^ and spatially intermediate between the extant diploid ancestors.

We also find no support for the Edger et al.^1^ hypothesis that the hexaploid *F. moschata* is a direct ancestor of the octoploids (Table 3, Figs. 4–5). Instead, its progenitors appear restricted to the “*vesca* clade” (*F. vesca, F. mandshurica, F. bucharica*).

Polyploidy is increasingly being recognized for its role in generating novel diversity in organismal evolution^10^, and the differential retention of diploid progenitors’ genes is central to this process. In particular, these diploid progenitors have each experienced a unique set of environments, pathogens and other challenges to survival, so prior to coming together in a polyploid genome, each progenitor independently evolved “answers” to a diverse array of abiotic and biotic conditions. Understanding the biology of these diploid species can inform polyploid plant adaptation to changing climate and associated environmental stress^11^. For these reasons, it is critical to correctly identify the diploid progenitors of polyploids.

## Materials and Methods

Illumina sequencing libraries were prepared from leaf tissue of six diploid *Fragaria* species, and 9-32X genomic coverage was obtained (Table 2). *Fragaria vesca* subsp. *bracteata, F. iinumae* and *F. nipponica* were sequenced at the Oregon State University Center for Genome Research and Biocomputing Central Services Lab with 100 bp single ends on a HiSeq 2000. *Fragaria mandshurica, F. nilgerrensis* and *F. nipponica* were sequenced at Berry Genomics (Hangzhou, China) with 150 bp paired ends on a HiSeq 2000. We removed adapters and low-quality portions of reads using Trimmomatic (v 0.35)^12^ and settings LEADING:20, TRAILING:20, SLIDINGWINDOW:5:20, MINLEN:50. The genome assemblies of octoploid *Fragaria ananassa* ‘Camarosa’ and diploid outgroup *Potentilla micrantha*^13^ were downloaded from the Genome Database for Rosaceae (https://www.rosaceae.org/). The former was subdivided by the Edger et al.^1^ based on diploid subgenome assignment into four sets of seven chromosomes. The outgroup and subgenome assemblies were converted to 20X genomic coverage of random 100 bp sequences using BBTools randomreads.sh (https://jgi.doe.gov/data-and-tools/bbtools/). The above sets of sequence reads were aligned with BWA (v 0.7.12)^14^ to the *Fragaria vesca* v 4.1 genome assembly^15^ after masking with the *F. vesca* v 4.1 transposable element library downloaded from the Genome Database for Rosaceae (https://www.rosaceae.org/).

The twelve resulting alignments were converted to a variant call format (vcf) file with SAMtools (v 1.9)^16^ with the default settings of the mpileup and call options. The vcf file was converted into a multisample variant format (mvf) file using MVFtools (v 0.5.1.4)^17^. All heterozygous sites were converted to N, to account for the fact that the octoploid subgenome and outgroup sequences were derived from haploid genome assemblies. MVFtools was used to automate maximum-likelihood (ML) estimates of phylogeny using RAxML (v 8.2.12)^18^ with the GTR+Γ model of sequence evolution and 100 bootstrap replicates. A taxon was excluded from the analysis of a 100 kb window if it had <10% of aligned sites. Analyses were conducted for the seven base chromosomes (Fig. 1b, Fig. 2) and for 2191 non-overlapping windows of 100 kb across the seven chromosomes (Fig. 1c, Fig. 3, Supplementary Table 1). The phylogenetic position of each subgenome relative to diploid species was recorded (see Fig. 3 for details) and summarized for each base chromosome (Supplementary Table 1). Homeologous exchange was inferred when the ‘Camarosa vesca’ subgenome shared a MRCA with *F. iinumae* or when the other three subgenomes shared a MRCA with the *F. vesca* clade (*F. vesca, F. mandshurica, F. bucharica*).

We used an F_1_ cross for linkage mapping of the hexaploid *F. moschata*. Our F_1_ mapping population was derived from two parental plants collected from Slovenia (46.6827°N, 16.2951°W). Following our previous protocols^8^, seeds of the experimental cross (*N* = 192) were planted in a custom soil mixture (2 : 1, Fafard 4 : sand top-dressed with Sunshine Redi-earth Plug & Seedling; Sun Gro Horticulture, Agawam, MA, USA), and grown under 16 °C /21 °C night/day temperatures and a 14-h photoperiod in a growth chamber at the University of Pittsburgh for 11 wk. We selected a random subset (*N* = 46) of the F_1_ progeny for targeted sequence capture.

Targeted sequence capture was performed using previously developed *Fragaria* baits (v 2.0)^8,19^. These 20,000 capture baits of 100 bp each are relatively randomly distributed across the seven base chromosomes (1–7). DNA was isolated from silica-dried leaf tissue of the 46 progeny and two parents at Ag-Biotech (Monterey, CA). We constructed individually indexed genomic libraries using the NEBNext Ultra DNA Library Prep Kit (New England BioLabs, Ipswich, MA, USA), which were then target enriched^8^ and sequenced using a 1/3 lane of 150 bp paired ends on a HiSeq 3000 at the Oregon State University Center for Genome Research and Biocomputing Central Services Lab.

The hexaploid linkage mapping involved four steps: quality filtering of paired-end capture reads, mapping reads to the above-mentioned diploid *F. vesca* v 4.1 genome assembly, genotype calling in polyploids using POLiMAPS^2^, and linkage mapping using OneMap^20^. First, we removed adapters and low-quality portions of paired-end reads using Trimmomatic (v 0.35)^12^ as described above, and merged the paired-end reads using PEAR (v 0.9.6)^21^ with a minimum overlap size of 20 bp. Second, we mapped both the merged and un-merged paired-end reads to the *F. vesca* v 4.1 reference using BWA (v 0.7.12)^14^. The sorted BAM files were generated with SAMtools (v 1.9)^16^ and then were used to create the mpileup file. Third, with the mpileup file, we conducted polyploid genotype calling using POLiMAPS^2^, in which heterozygous and homozygous loci were identified with the default parameters, except for the depth of ≥32X per progeny for the hexaploid. Lastly, to construct maternal and paternal linkage groups (LGs), we assigned SNPs to the most likely LGs based on a logarithm of odds (LOD) threshold of 5 using OneMap^20^ in R (v 3.3.3)^22^. LGs with at least 20 SNPs were used for subsequent analyses.

To infer the phylogenetic replacement of *F. moschata* in relation to the octoploid strawberry and other diploid *Fragaria*, we first extracted the quality-filtered reads that contained the above LG SNPs, and mapped these reads to the *F. vesca* v 4.1 reference to generate consensus LG sequences using POLiMAPS. We next used POLiMAPS and the variant call format file described in the above Phylogenomic Analysis of Octoploid Subgenomes section to generate multiple sequence alignments among *F. moschata* LG sequences, the four octoploid reference subgenomes, diploid *Fragaria* genomes and the outgroup *Potentilla*. Phylogenetic inference was conducted for each of chromosomes 1–7 and for maternal and paternal LGs separately, using the ML method with the GTR+Γ model and 100 bootstrap replicates in RAxML (v 8.0.26)^18^.

To test the hypothesis of Edger et al.^1^ that *F. nipponica* and *F. viridis* are the progenitors (sister taxa) of the respective ‘Camarosa_nipponica’ and ‘Camarosa_viridis’ subgenomes, we built the ML tree of constrained topology that reflects this hypothesis ((Camarosa_nipponica, nipponica),(Camarosa_viridis, viridis),(Camarosa_vesca,bracteata),(Camarosa_iinumae, iinumae)) (Table 3), for each of the seven base chromosomes. These constrained trees were then compared to the ML unconstrained trees using the Shimodaira–Hasegawa (SH) test in RAxML.

To test the hypothesis of Edger et al.^1^ that the hexaploid *F. moschata* is an evolutionary intermediate in the formation of the octoploid strawberry, we built ML trees of constrained topologies that reflect this hypothesis (Table 4). Specifically, under this hypothesis, each of the three maternal or paternal LGs should be sister to one of the octoploid subgenomes (i.e. Camarosa_viridis, Camarosa_iinumae, and Camarosa_nipponica). We built six constraint trees by alternating the combinations of the three LGs and the three octoploid subgenomes, for each of chromosomes 1–7 and for maternal and paternal LGs separately (Table 4). These constrained trees were then compared to the ML unconstrained trees using the SH test in RAxML.

## Acknowledgements

The authors thank Lucas Longway and Rich Cronn for laboratory assistance, Kevin Weitemier and Tara Jennings for assistance with targeted sequence capture, and Katherine Schuller, Elizabeth Jiang, Anna Freundlich and the University of Pittsburgh glasshouse staff for plant cultivation. The OSU Center for Genome Research and Biocomputing conducted Illumina sequencing and provided computational support. This work was funded by the US National Science Foundation (DEB 1241217 to A.L. & DEB 1020523 to T.L.A.) and the National Natural Science Foundation of China (No. 31261120580 to M.D. and J.L.).

## Author Contributions

*Fragaria moschata* linkage mapping was conducted by N.W. and T.L.A. Phylogenetic analyses were performed by A.L., N.W., and J.T. Figures were prepared by J.T. and N.W. Sample collection and DNA sequencing in China was led by J.L. and M.D. All authors contributed to the writing and reviewing of the manuscript.

## Competing Interests

The authors declare no competing interests.

## Data Availability

The sequence alignments and phylogenetic trees will be available on Dryad. The raw sequence data will be available in the Sequence Read Archive under an NCBI BioProject.

